# How Much Does TRPV1 Deviate from An Ideal MWC-Type Protein

**DOI:** 10.1101/2023.10.26.564268

**Authors:** Shisheng Li, Jie Zheng

**Author notes:** Correspondence should be sent to: Jie Zheng. **Author contributions:** Conceptualization: SL, JZ; Patch clamp: SL; Data analysis: SL, JZ; Funding acquisition: JZ; Supervision: JZ; Writing – original draft: SL, JZ; Writing – review & editing: SL, JZ.

## Abstract

Many ion channels are known to behave as an allosteric protein, coupling environmental stimuli captured by specialized sensing domains to the opening of a central pore. The classic Monod-Wyman-Changeux (MWC) model, originally proposed to describe binding of gas molecules to hemoglobin, has been widely used for analyzing ion channel gating. Here we address the issue of how accurate the MWC model predicts activation of the capsaicin receptor TRPV1 by vanilloids. Taking advantage of a concatemeric design that makes it possible to lock TRPV1 in states with zero-to-four bound vanilloid molecules, we showed quantitatively that the overall gating behavior is satisfactorily predicted by the MWC model. There is however a small yet detectable subunit position effect: ligand binding to two kitty-corner subunits is 0.4-to-0.6 kcal/mol more effective in inducing opening than binding to two neighbor subunits. This difference, less than 10% of the overall energetic contribution from ligand binding, is not expected in hemoglobin, in which each subunit is related equivalently to all the other subunits.

**Significance:** The MWC model, proposed more than 50 years ago, is elegantly simple yet powerful in predicting the behavior of allosteric proteins like hemoglobin. Its prediction power for ion channel gating has been beautifully demonstrated in the studies of BK channels. Our present work aims to determine how accurate the MWC model predicts TRPV1 activation induced by vanilloids. Our findings support the notion that the evolutionary drive upon allosteric proteins applies generally to multi-subunit proteins.

## Introduction

In the landmark study of neuronal action potential, Hodgkin and Huxley found that the entities controlling transmembrane conductance for sodium and potassium ions—which we now know as voltage-gated sodium (Nav) channels and voltage-gated potassium (Kv) channels—operate with high voltage sensitivities (Hodgkin and Huxley 1952). In their empirical equations describing the voltage dependence of sodium and potassium conductance, G_Na_ and G_K_, respectively, this high sensitivity is reflected by the exponents assigned to the probability terms. Modern expansions of the H&H ideas incorporating knowledge of Nav and Kv channel structures reveal that high voltage sensitivity is partially rooted in the highly cooperative nature of voltage-dependent activation. In the 1990s, Zagotta, Hoshi and Aldrich (Zagotta, Hoshi, and Aldrich 1994) and Schoppa and Sigworth (Schoppa and Sigworth 1998) identified a late cooperative transition that would be needed in an otherwise independent subunit gating scheme to satisfactorily describe the course of activation in *Shaker* potassium channels. It is thought that the voltage sensors of *Shaker*, as well as many other voltage-gated ion channels, operate in a mostly independent manner, whereas some of the conformational changes in the channel pore must be concerted (Zheng and Trudeau 2023).

BK potassium channels are activated by not just voltage but also intracellular calcium. Their activation also exhibits cooperative features (Rothberg and Magleby 1998). In a comprehensive investigation of BK channel macroscopic currents, single-channel currents, and gating currents, Horrigan, Cui and Aldrich revealed that the voltage sensor and calcium sensor operate separately, and they both influence the channel pore opening allosterically (Horrigan and Aldrich 1999; Horrigan, Cui, and Aldrich 1999; Horrigan and Aldrich 2002). The kinetic model that could satisfactorily describe BK activation behaviors thus contains two branches of allosteric coupling. In each branch of the model, the four sensors (for voltage or calcium) contribute an identical amount of energy towards influencing the pore opening.

This type of allosteric coupling has been previously proposed to govern another protein, the hemoglobin. Hemoglobin is a symmetrical tetramer, with each of the four subunits contains a binding pocket for gas molecules (Bolton and Perutz 1970). Building upon previous findings that oxygen binding to hemoglobin is cooperative (Edsall 1972), Monod, Wyman and Changeux proposed that the cooperativity comes from a concerted conformational change that affects all gas binding sites equally (Monod, Wyman, and Changeux 1965). Hemoglobin serves as a carrier for gas molecules; there is no function equivalent to ion conduction in an ion channel that can be used as a direct indicator of the concerted transition. Studies of hemoglobin therefore focused on the ligand binding process. It was proposed that, there is an equal energetic contribution to the concerted transition by each gas molecule binding step, and these binding steps are independent (Monod, Wyman, and Changeux 1965). Introducing interactions at the binding steps, such as those seen in the sequential model, yields good performance but also adds complexity (Koshland, Némethy, and Filmer 1966).

Equal and independent contribution to gating by each subunit is assumed for BK channels in the study by Horrigan, Cui and Aldrich, and for many other ion channels studied subsequently. The MWC-type models in general worked well in predicting channel behaviors and provided important guidance for mechanistic investigations in the following decades. Many predictions from these studies are nicely confirmed when ion channel structures became available (Zheng and Trudeau 2023). In recent studies of the capsaicin receptor TRPV1 (Li et al. 2023; Li and Zheng 2023), we realized that a set of concatemers previously designed by Priel and colleagues (Hazan et al. 2015) would allow us to lock a TRPV1 channel in each of the intermediate ligand-bound states, hence presenting a unique opportunity to directly test the applicability of the MWC model to the TRPV1 channel. This is possible because the concatemers, made of various combinations of wildtype and Y511A mutant protomers, would trap a vanilloid molecule such as resiniferatoxin (RTX) when it binds to a wildtype subunit but allow it to fall off a mutant subunit. Isolating intermediate binding states for equilibrium measurements has been challenging for hemoglobin bound with small gas molecules; it is, to our best knowledge, not done with any concatemeric ion channels. In the present study, we took advantage of this powerful system to further evaluate TRPV1 activation by capsaicin.

## Materials and Methods

### Molecular Biology

The plasmids used in this study were made in the Priel laboratory. Briefly, wild-type rat TRPV1 (Y) cDNAs were joined with the Y511A mutant (A) cDNAs in various combinations: YYYY, YYYA, YYAA, YAYA, AYAY, YAAA, and AAAA. Their functional properties have been carefully tested and described in previous publications (Hazan et al. 2015; Li et al. 2023). Representative single-channel recordings are presented in Supplementary Figures.

### Cell Culture

TSA201 cells (HEK293T variant from American Type Culture Collection) served as the expression system for patch clamp assays. These cells were cultivated on 25mm glass coverslips in 30mm dishes (from Fisher Scientific) until reaching 30% - 50% confluence, then transiently transfected. The transfection was carried out using Lipofectamine 2000 (Invitrogen) 24 hours prior to patch-clamp recording, following the manufacturer’s guidelines. For single-channel recordings, a combination of 0.1 μg concatemer plasmid and 0.2 μg eYFP plasmid was utilized per transfection.

### Chemical Solutions

For inside-out patch-clamp recordings, we used symmetric bath and pipette solutions containing 140 mM NaCl, 15 mM HEPES, 2 mM EDTA, with a pH of 7.4. Capsaicin (from Sigma-Aldrich) was dissolved in Dimethyl sulfoxide (DMSO) to prepare a 1 M stock, then further diluted to concentrations ranging from 0.01 μM to 100 μM using the bath solution. Both resiniferatoxin (Alomone Labs) and 6’-iodoresiniferatoxin (6’-iRTX, Sigma-Aldrich) were dissolved in ethanol to create a 1 mM stock and subsequently diluted to a 200 nM working concentration with the bath solution. Lastly, 2-Aminoethoxydiphenyl borate (2-APB, Sigma-Aldrich) was dissolved in DMSO to produce a 1 M stock, which was then diluted to a 3 mM working solution.

### Electrophysiology

Pipettes for patch-clamp recordings were pulled from borosilicate glass capillaries (Sutter Instrument) using a P-97 micropipette puller (Sutter Instrument) and fire-polished to achieve resistances of 2-6 MΩ for macroscopic recordings and 8-15 MΩ for single-channel recordings. We employed an EPC 10 USB patch-clamp amplifier (Warner Instruments), operated by the PatchMaster software. Sampling and filtering frequencies were set at 10 kHz and 2.25 kHz, respectively. Patch clamp configurations were primarily inside-out, unless specifically stated otherwise. The holding potential began at 0 mV and proceeded to steps of +80 mV and –80 mV. Step durations were adjusted as required. A gravity-driven perfusion system, managed by the Rapid Solution Changer (RSC200, Biological Science Instrument), facilitated solution perfusion and changes.

### Data Analysis

Patch-clamp data, exported from PatchMaster in the Igor format, were analyzed using Igor Pro 8 (WaveMetrics). Statistical analyses were conducted in Graphpad Prism 8, employing either the Student t-test or one-way ANOVA based on the specific requirements.

Open probabilities of single channels were determined by creating all-point histograms from single-channel current traces. The open probability was calculated as 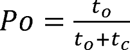, where *t*_0_ denotes the total channel open time and *t_c_* the total channel closed time. For patches containing two channels, open probability was derived using: 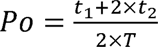. Here, *t*_l_ represents the time with a single channel opening, *t*_2_ the time with both channels opening, and *T* the overall analyzed duration. Recordings with more than two channels were discarded.

### Model fitting

The global fitting package in Igor Pro 8 was used to fit single channel Po data from the concatemers. To estimate the binding effect, the following equation was used:

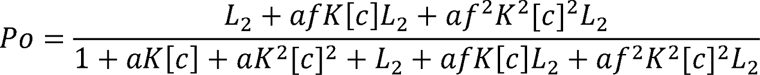

where *L*_2_ is the equilibrium constant for channels pre-loaded with two RTX or 6’-iRTX molecules, [*c*] is the concentration of capsaicin in the perfusion solution, *K* is the binding affinity parameter, *a* is the cooperativity coefficient for binding (ratio between the two binding affinity parameter generated by first and second capsaicin binding), f is the gating parameter (fold change that each capsaicin bound would shift the equilibrium from closed to open state).

To estimate the gating effect, the following equation was used:

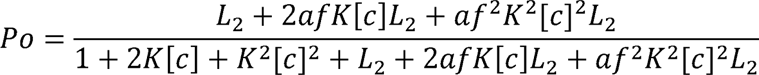

where *L*_2_ is the equilibrium constant for channels was pre-loaded with two RTX or 6’-iRTX molecules, [*c*] is the concentration of capsaicin in perfusion solution, *K* is the binding affinity parameter, *f* is the gating parameter (fold change that each capsaicin bound would shift the equilibrium from closed to open state) and *a* is the cooperativity coefficient for gating (ratio between the two gating parameter generated by first and second capsaicin binding).

To directly determine the energy contribution from gating by measuring the Po with different number of bound RTX molecules, the following equations were used:

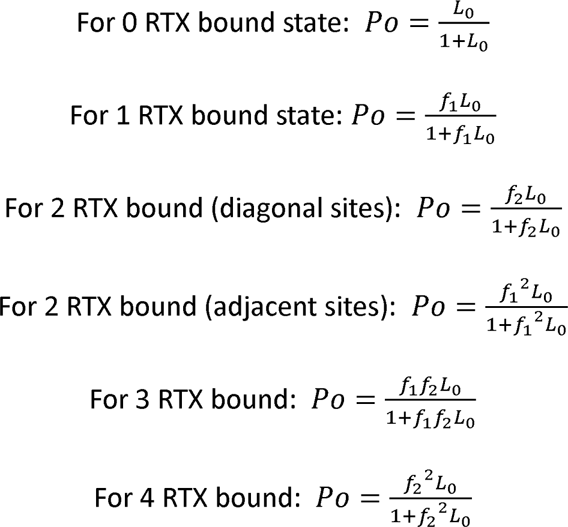

Where *L*_O_ is the equilibrium constant for channels in the apo state, *f*_l_ is the gating parameter when only one RTX bound in the diagonal pair of the binding sites, *f*_2_ is the gating parameter when two RTX bound in the diagonal pair of the binding sites.

## Results

TRPV1 is an allosteric protein. In the absence of ligands (such as capsaicin or RTX), TRPV1 can spontaneously transition from a closed state to an open state at a very low probability (with an equilibrium constant of less than 0.01) (Li et al. 2023; Yang et al. 2018). As an increasing number of agonist molecules bind to TRPV1, the equilibrium progressively shifts towards the open state. Our previous study showed that, during RTX activation of TRPV1, sequential RTX bindings shift the equilibrium nearly exponentially, suggesting the energy contributions from these RTX binding steps are approximately equal (Li et al. 2023), as the classic MWC model postulates (Monod, Wyman, and Changeux 1965). Binding of all four subunits with RTX molecules contributes a total energy of 6.8-to-7.4 kcal/mol towards activation (Li et al. 2023).

However, when we closely examined the capsaicin response curves for YAYA, AYAY, and YYAA concatemers—all producing channels with two wild-type subunits and two mutant subunits but with different subunit arrangements (Figure 1A)—a slight but apparent difference could be discerned. Both YAYA and AYAY (with two mutant subunits located at kitty-corners in the assembled channels) appeared to be slightly more sensitive to capsaicin than YYAA (with mutant subunits at neighbor positions) (Figure 1B). The small differences in the EC50 value, obtained from fitting with a Hill function, would translate into an energy difference of about 0.66 kcal/mol between YYAA and YAYA, and 0.53 kcal/mol between YYAA and AYAY. Nonetheless, at most capsaicin concentrations, a statistically significant difference in Po was hard to be confidently established .

**Figure 1.**
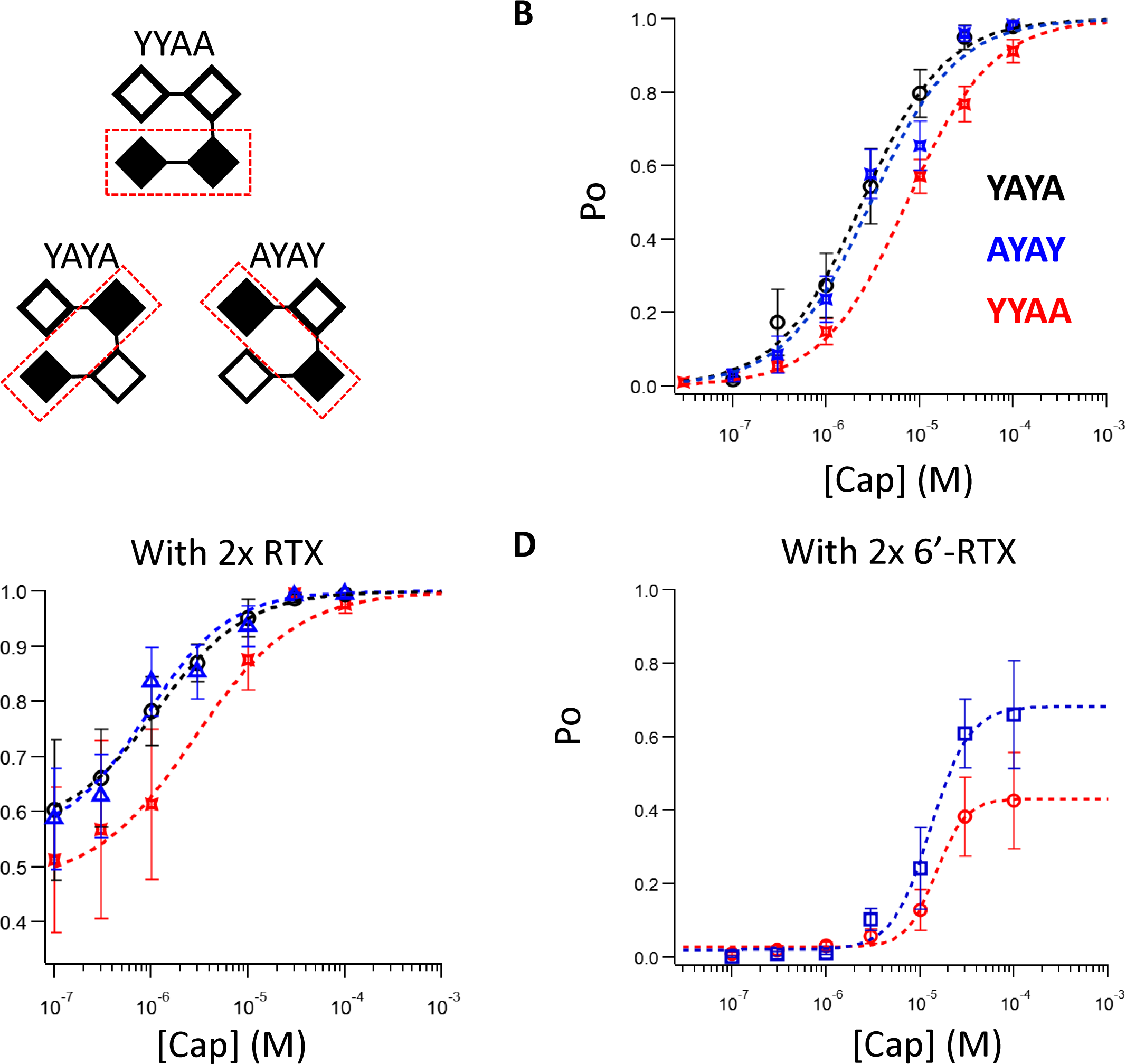
A. Illustration of the design of YYAA, YAYA and AYAY concatemers; open square represents a wild type subunit; filled square represents a Y511A mutant subunit. B. Capsaicin concentration-dependent TRPV1 single-channel open probability curves for YYAA, YAYA and AYAY fitted to a Hill Equation using the following EC50 and Hill slope parameters: YYAA (red), 7.31 μM, 0.87, n = 12; AYAY (blue), 2.99 μM, 0.95, n = 4; YAYA (black), 2.42 μM, 0.97, n = 5. C. Capsaicin concentration-dependent TRPV1 single-channel open probability curve for YYAA, YAYA and AYAY with two preloaded RTX molecules in the wildtype protomers fitted to a Hill Equation: YYAA, 2.81 μM, 0.82, n = 5; AYAY, 0.90 μM, 1.02, n = 5; YAYA, 1.14 μM, 0.94, n = 5. D. Capsaicin concentration-dependent TRPV1 single-channel open probability curve for YYAA and AYAY with two preloaded 6’-iRTX molecules in the wildtype protomers fitted to a Hill Equation: YYAA, 14.8 μM, 2.72, n = 4; AYAY, 13.0 μM, 2.11, n = 4. Data are presented as mean ± SEM.

Given the uncertainty, we assessed the potential subunit positioning effect with a different approach. Taking advantage of the reversible binding of RTX to the A subunits and irreversible binding to the Y subunits, we first fully loaded each channel type with RTX, followed by thorough washing. As we reported previously (Li et al. 2023; Li and Zheng 2023), this procedure yielded channels containing two RTX molecules that bound to the Y subunits and two A subunits available for subsequent binding by capsaicin. We measured the channel open probability at increasing concentrations of capsaicin, which was normalized to that when the channel was fully loaded with four RTX molecules. As shown in Figure 1C, a slightly higher capsaicin sensitivity in YAYA and AYAY concatemeric channels was again observed, which is reflected in the uplifted concentration-dependence curves, though at most capsaicin concentrations this apparent difference wasn’t statistically significant. Similar results were obtained using 6’-iRTX, a RTX derivative that is a much weaker TRPV1 agonist; an upward shift in the AYAY curve could be seen (Figure 1D). (The YAYA concatemer exhibited unusually high Po values under this experimental condition and was excluded from this and subsequent analyses.) Hill fitting indicated that the left-shifted EC50 values for YAYA and AYAY were equivalent to an energy difference of 0.53-0.68 kcal/mol, without a substantial change in the Hill slope. Therefore, we hypothesized that there is a detectable difference in activation energy for ligand binding to diagonal versus adjacent subunits.

Our results indicate that, when two ligands (RTX or 6’-iRTX) bind to a single TRPV1 channel, the diagonal (“kitty-corner”) binding positions produce a higher open probability than the adjacent binding positions. This behavior is not aligned with the prediction of the classic MWC model (Figure 2A). To experimentally assess the magnitude of this position effect, we used a modified model incorporating distinct states for the two-ligand bound configurations, as shown in Figure 2B. In this expanded model, the increased cooperativity for the diagonal binding positions could arise either from the binding process (characterized by the K parameter) or the gating process (represented by the f parameter). We evaluated these possibilities separately in the following two sets of experiments, using single-channel Po data in response to capsaicin after the two Y subunits in each channel were preloaded with RTX or 6’-iRTX molecules, as described earlier.

**Figure 2.**
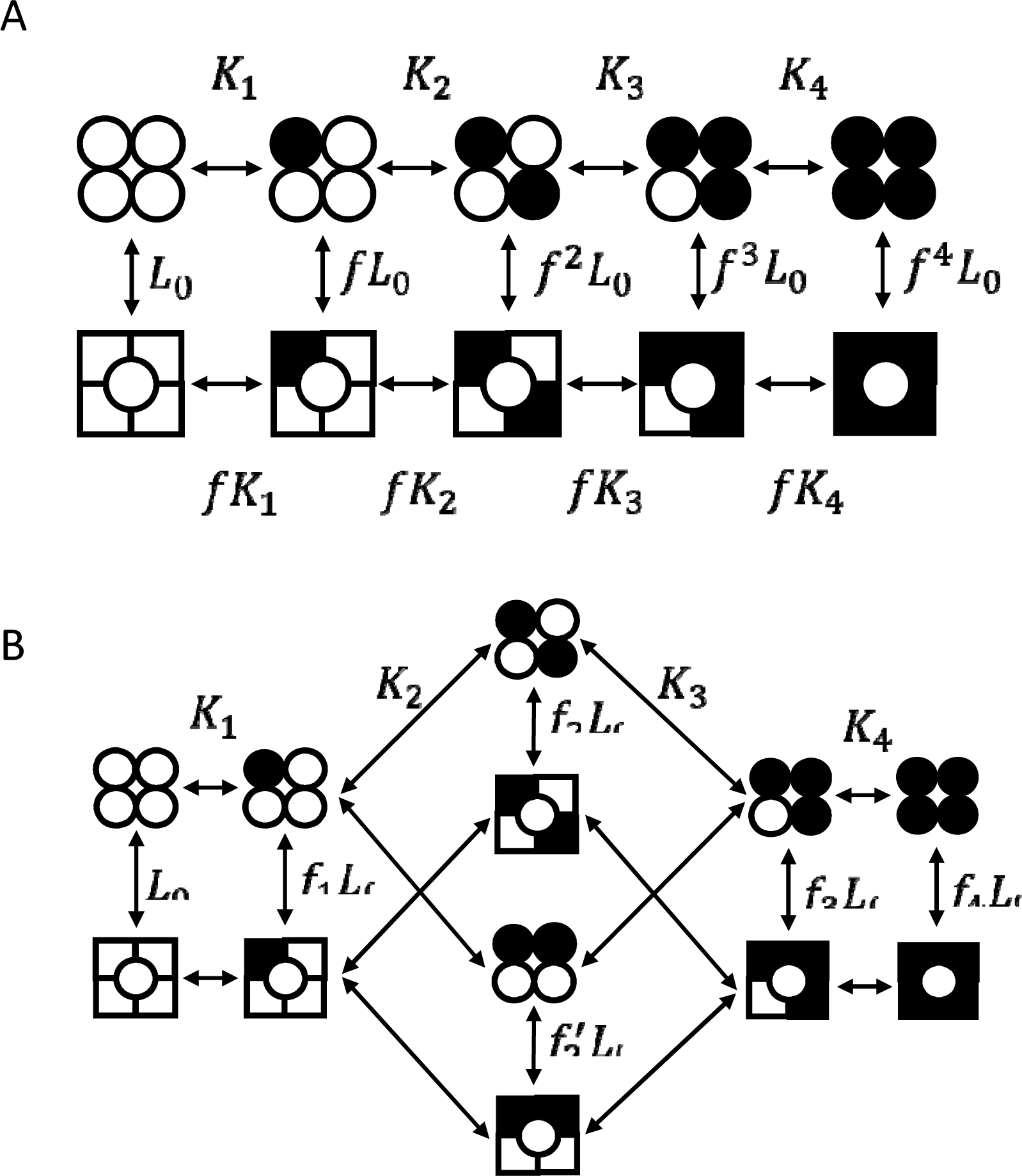
A. Illustration of a classic MWC model for a tetrameric ligand-gated ion channel with four identical ligand-binding sites. L_0_ represents the equilibrium constant for the apo state, f represents the cooperative factor which reflects the energy contribution by each ligand binding; K_1_ to K_4_ represent the ligand association constants. B. Illustration of a modified model with positional effects.

To assess a potential subunit positional effect on binding energy, we first assumed the gating effect (f) to be independent of the positional effect (Figure 3A&B); that is, variation in gating cooperativity was assumed to originate solely from binding, hence potentially exaggerating the binding effect. We assigned the equilibrium constant for the last binding step the same K parameter, since this step is identical for all concatemers, as can be seen in Figure 3A&B. However, transitioning from a two-ligand-bound state to a three-ligand-bound state might show sensitivity to the initial positions of liganded subunits, leading us to introduce a coefficient factor a or a’ to the binding parameter K. In the absence of a positional effect, both a and a’ would be 4 as expected from independent binding. However, if a positional effect existed, a and a’ would differ. Global fitting of datasets from all concatemers pretreated with either RTX or 6’-iRTX (Figure 3C) yielded values of a = 3.03 and a’ = 4.33, suggesting a positional effect, albeit one that is rather small. The calculated free energy difference between the two binding configurations, derived from the ratio of a’/a, stands at 0.21 kcal/mol. Considering that our method likely overestimated binding cooperativity, the actual difference would be smaller. An RMSE of 0.033 for the global fitting suggests that our model adequately represents channel behavior.

**Figure 3.**
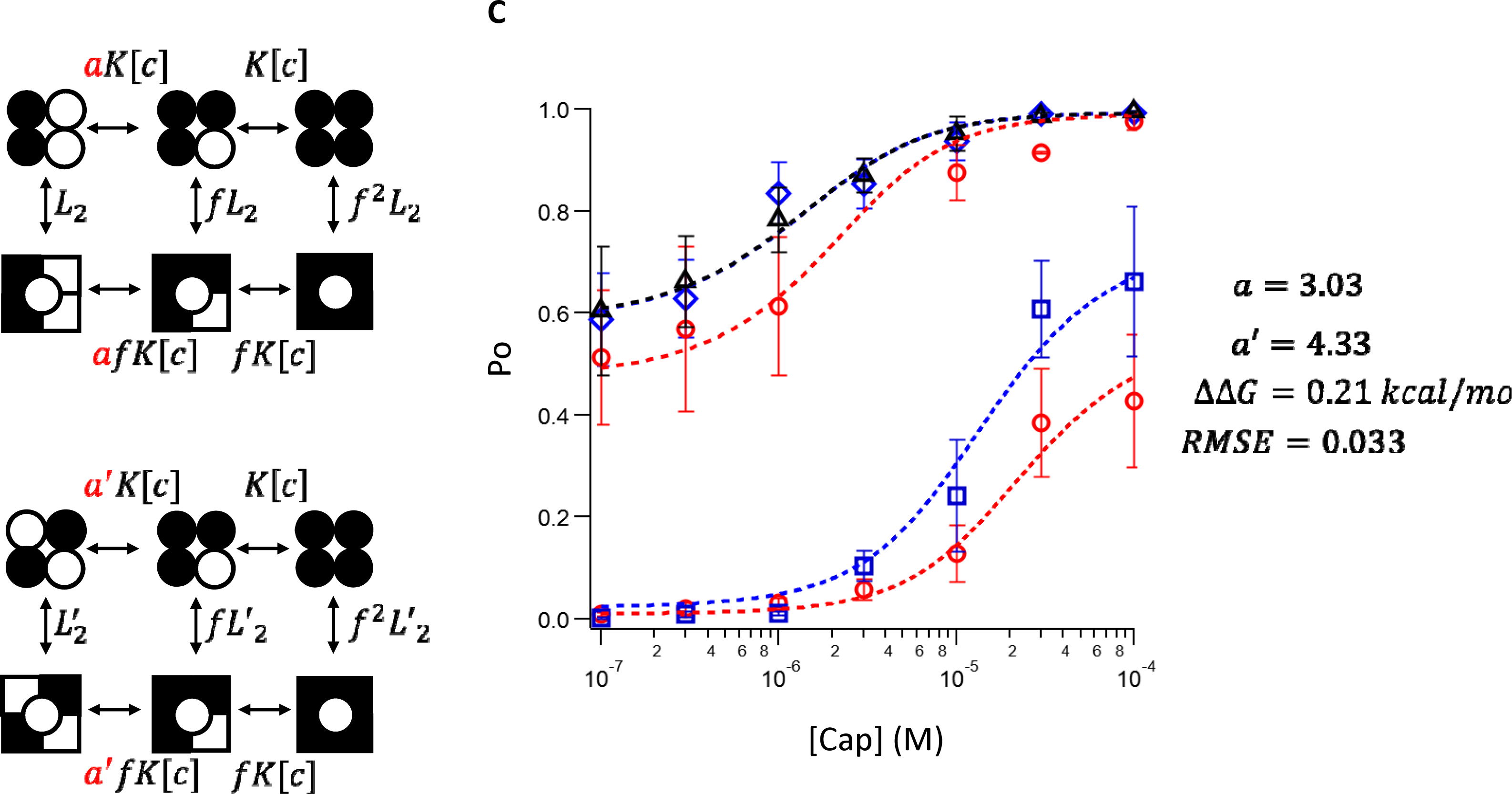
A&B. Illustrations of the model used for fitting the capsaicin-dependent Po data with two pre-loaded RTX or 6’-iRTX, to test the contribution of binding to the positional effect. L_2_ and L_2_’ represent the equilibrium constant for the state with two preloaded RTX or 6’-iRTX (L_2_ for YYAA, L_2_’ for YAYA and AYAY), before application of capsaicin; a and a’ are coefficient factors for the positional effect (a for YYAA, a’ for YAYA and AYAY); when there is no positional effect, a and a’ should be 4 for independent bindings; f represents the cooperative factor which reflects the energy contribution by each ligand binding; K represents the ligand binding affinity constant for the Y511A mutant subunit, [c] is the capsaicin concentration. C: Global fitting results using the model from panels A and B with the following parameters: a = 3.03; a’ = 4.33. These fittings yield ΔΔ*G* = 0.21 kcal/mol, and RMSE = 0.033.

To gauge the gating effect, we used a similar approach, assuming the binding steps were independent (Figure 4A&B). This would attribute any binding cooperativity to gating, potentially exaggerating the gating effect. For the reason discussed earlier, we assigned the same f factor when the channel transitioned from a three-ligand-bound state to a four-ligand-bound state. We introduced a coefficient factor a or a’ to the gating parameter f for transitions from a two-ligand-bound state to a three-ligand-bound state. Without a positional effect, a and a’ would be 1; otherwise, a and a’ would differ. Global fitting of the same datasets yielded values of a = 0.77 and a’ = 2.26. The free energy difference between these configurations, derived from a’/a, is 0.64 kcal/mol. Although our approach might exaggerate the gating effect, the larger energy difference clearly suggests that gating is more influential in the positional effect. The reduced RMSE of 0.030 indicates slightly improved model fitting.

**Figure 4.**
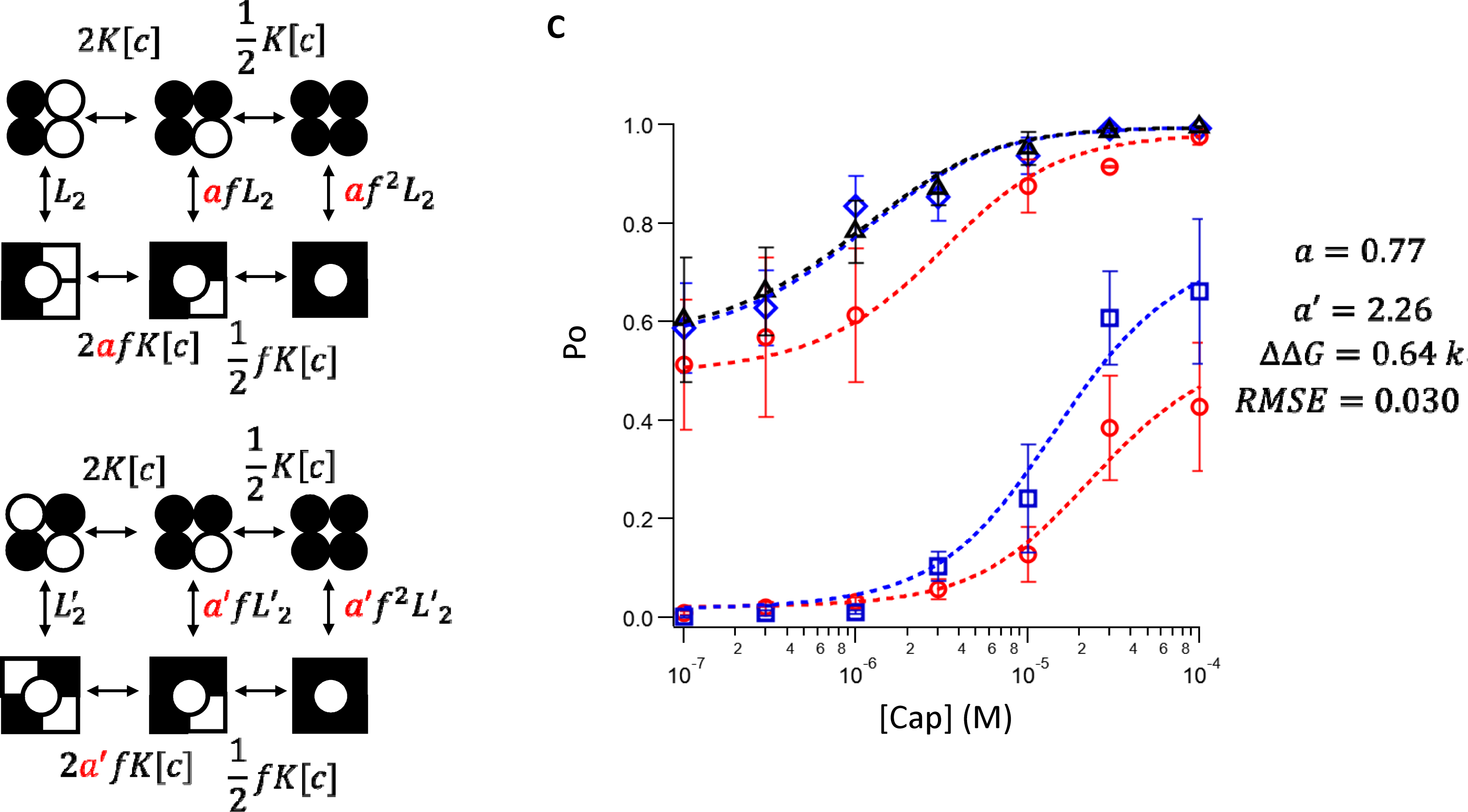
A&B. Illustrations of the model testing the contribution of gating to the positional effect. L_2_ and L_2_’ represent the equilibrium constant for the state with two preloaded RTX or 6’-iRTX (L_2_ for YYAA, L_2_’ for YAYA and AYAY), before application of capsaicin; a and a’ are coefficient factors for the positional effect (a for YYAA, a’ for YAYA and AYAY); when there is no positional effect, a and a’ should be 1; f represents the cooperative factor which reflects the energy contribution by each ligand binding; K represents the ligand binding affinity constant for the Y511A mutant subunit, [c] is the capsaicin concentration. C: Global fitting results using the model from panels A and B with the following parameters: a = 0.77; a’ = 2.26. These fittings yield ΔΔ*G* = 0.64 kcal/mol, and RMSE = 0.030.

Since both fitting methods could potentially skew the positional effect, we explored a direct approach. We have previously shown that, by fully loading each concatemer with RTX followed by thorough washing, we could isolate channels with one-to-four RTX-bound subunits (Li et al. 2023). Po measurements from these channels reflect only the gating equilibrium of each vertical transition seen in Figure 5A without confounding effects from binding. The difference between two-ligand-bound channels could be represented by introducing just one additional degree of freedom to the classic MWC model. We assigned f_1_ as the gating coefficient for a single independent binding site and hypothesized that each pair of diagonal binding sites would exhibit stronger cooperativity (f_2_) than two independent sites (f_1_^2^). Using this model to fit the RTX-bound TRPV1 single-channel Po data (Figure 5B), we found f_2_ to be 107% larger than f_1_^2^, equivalent to a free energy difference of 0.43 kcal/mol. This result aligns nicely with our indirect estimates discussed earlier, suggesting that while binding might contribute slightly to the positional effect (probably much less than 0.21 kcal/mol), gating plays a significant role, contributing approximately 0.43-to-0.64 kcal/mol. Compared to the 6.8-to-7.4 kcal/mol total energetic contribution from ligand binding to activation, this different is small (less than 10%).

**Figure 5.**
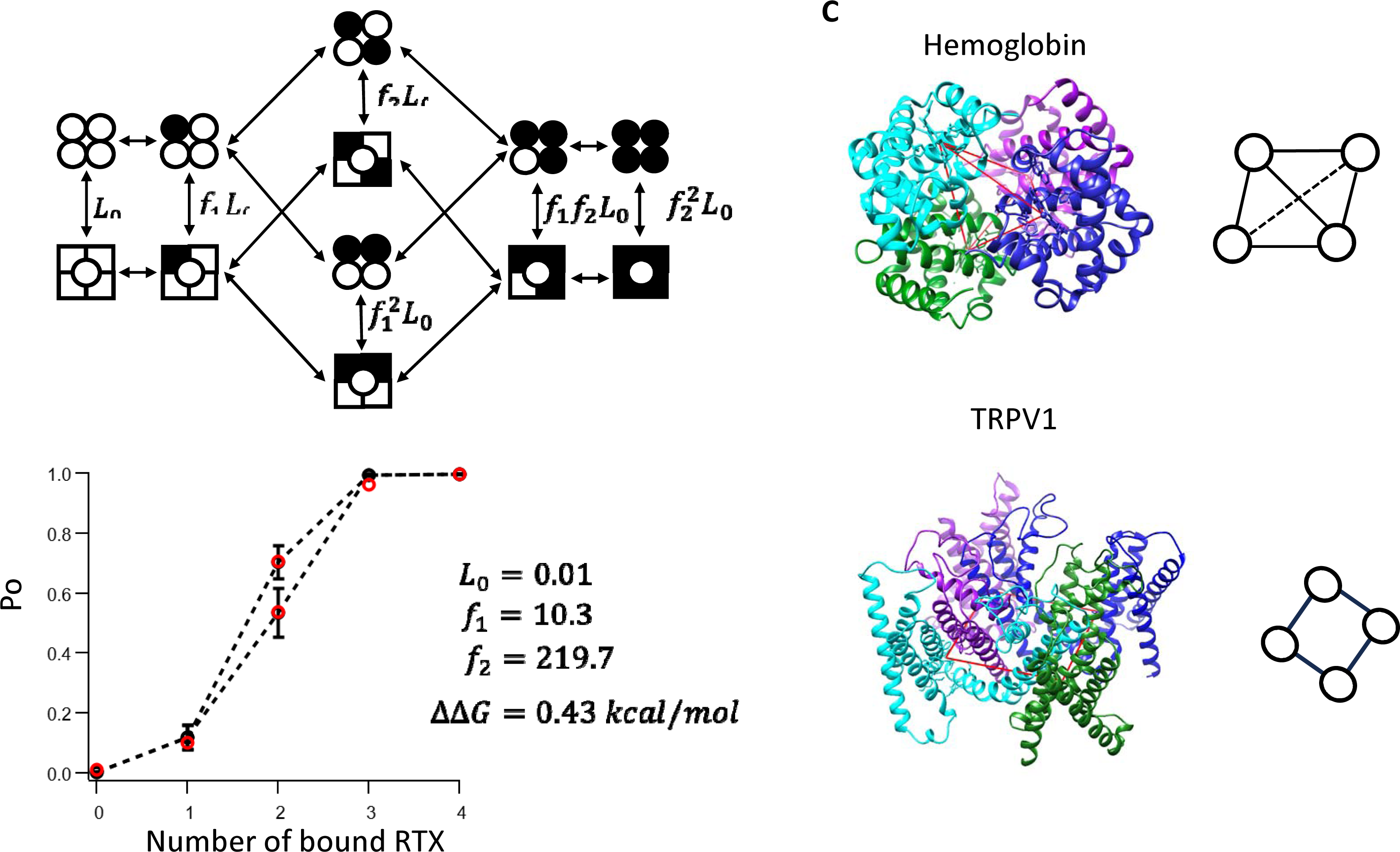
A. Illustration of a modified MWC model with positional effects. L_0_ represents the equilibrium constant for the apo state, f_1_ represents the cooperative factor which reflects the energy contribution by one ligand binding in the pair of the two diagonal binding sites; f_2_ represents the cooperative factor which reflects the energy contribution by two ligands binding to the pair of the two diagonal binding sites; when there is no positional effect, f_2_ should equal to f_1_^2^, making the model equivalent to the classic MWC model. B. Global fitting results for the Po data from AAAA, YAAA, YYAA, YAYA, AYAY, YYYA, YYYY concatemers loaded with a varying number of RTX with the following parameters: L_0_ = 0.01, f_1_ = 10.3, f_2_ = 219.7, which produce ΔΔ*G* = 0.43 kcal/mol, n = 4-17. Data are presented as mean ± SEM. C. Relationships between ligand binding pockets in hemoglobin (PDB entry 2DHB, top panel) and tetrameric ligand-gated ion channels such as TRPV1 (transmembrane part of TRPV1, modified from PDB entry 3J5R, bottom panel). Positions of bound ligands are highlighted by red lines between them, and illustrated by a diagram on the right.

## Discussion

The present paper is the third in a series of studies of TRPV1 activation by vanilloid molecules (Li et al. 2023; Li and Zheng 2023). Our collective results from these studies suggest that, as an allosteric protein, TRPV1 can exist in various conformations (closed versus open, and with various number of ligands) at any particular ligand concentration. A recent structural study indeed captured many of these conformations, including those with two bound ligands at either neighbor or kitty-corner positions (Zhang, Julius, and Cheng 2021). It remains unclear what might contribute to the small difference in stability between these two configurations. The cryo-EM structures nonetheless reveal that a vanilloid molecule bound in the vanilloid binding pocket does not belong strictly to one subunit: due to the domain-swapped arrangement, the vanilloid binding pocket is formed by the S3 and S4 segments and the S4-S5 linker from one subunit together with the S5 and S6 segments from a neighbor subunit (Liao et al. 2013). Whereas capsaicin forms hydrogen bonds with the S4 segment and the S4-S5 linker of the same subunit, extensive hydrophobic interactions with the S5 and S6 segments are predicted from both the cryo-EM structures and computational modeling (Yang et al. 2015). Polar interactions between capsaicin or other ligands and the S6 segment are also predicted (Yin et al. 2019; Vu et al. 2020; Dong et al. 2019).

One intriguing question arises from the present study is, does hemoglobin exhibit positional differences in ligand induced allosteric transition like what we have observed in TRPV1? To our best knowledge, no direct experimental evidence in support of such a possibility has been reported. The binding pockets for gas molecules in hemoglobin are formed within each of the four subunits as an isolated domain (Bolton and Perutz 1970). These binding pockets are positioned at the vertex corners of a tetrahedron, such that each pocket is related to all the other pockets nearly equally (Figure 5C, top panel). TRPV1 and other ion channels existing in a planar membrane do not share this symmetry. A vanilloid binding pocket in TRPV1 is related differently to its neighbor pockets and the kitty-corner pocket (Figure 5C, bottom panel). When two neighbor binding pockets are occupied by ligands, the channel complex lacks the rational symmetry exhibited when the kitty-corner binding pockets are occupied. Despite this geometric restriction imposed by the membrane, cooperative control of a centrally located ion permeation pore by its surrounding subunits bestows great sensitivities to the multi-subunit TRPV1 channel and most biological ion channels.

**Supplementary Figure 1a-b.**
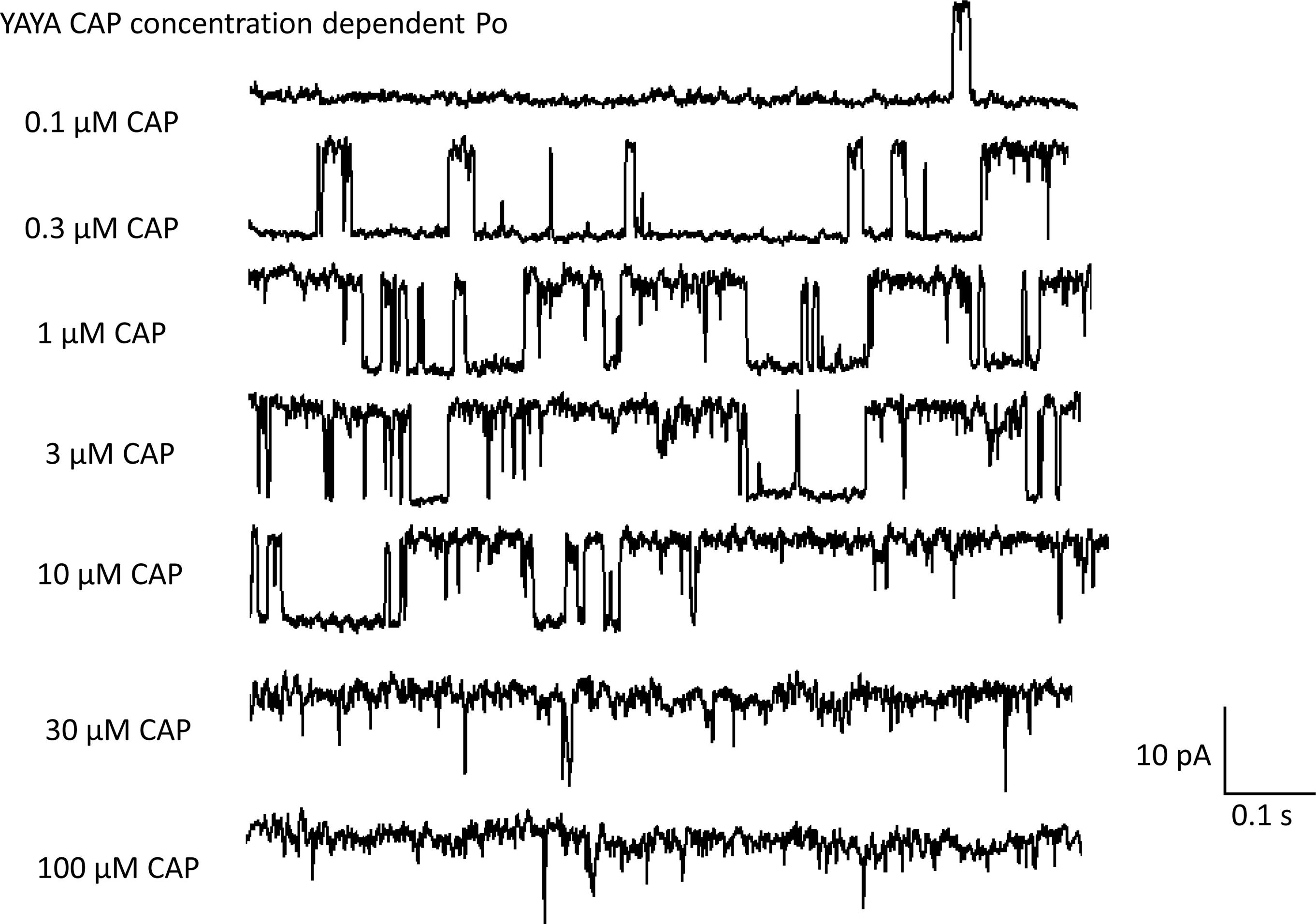
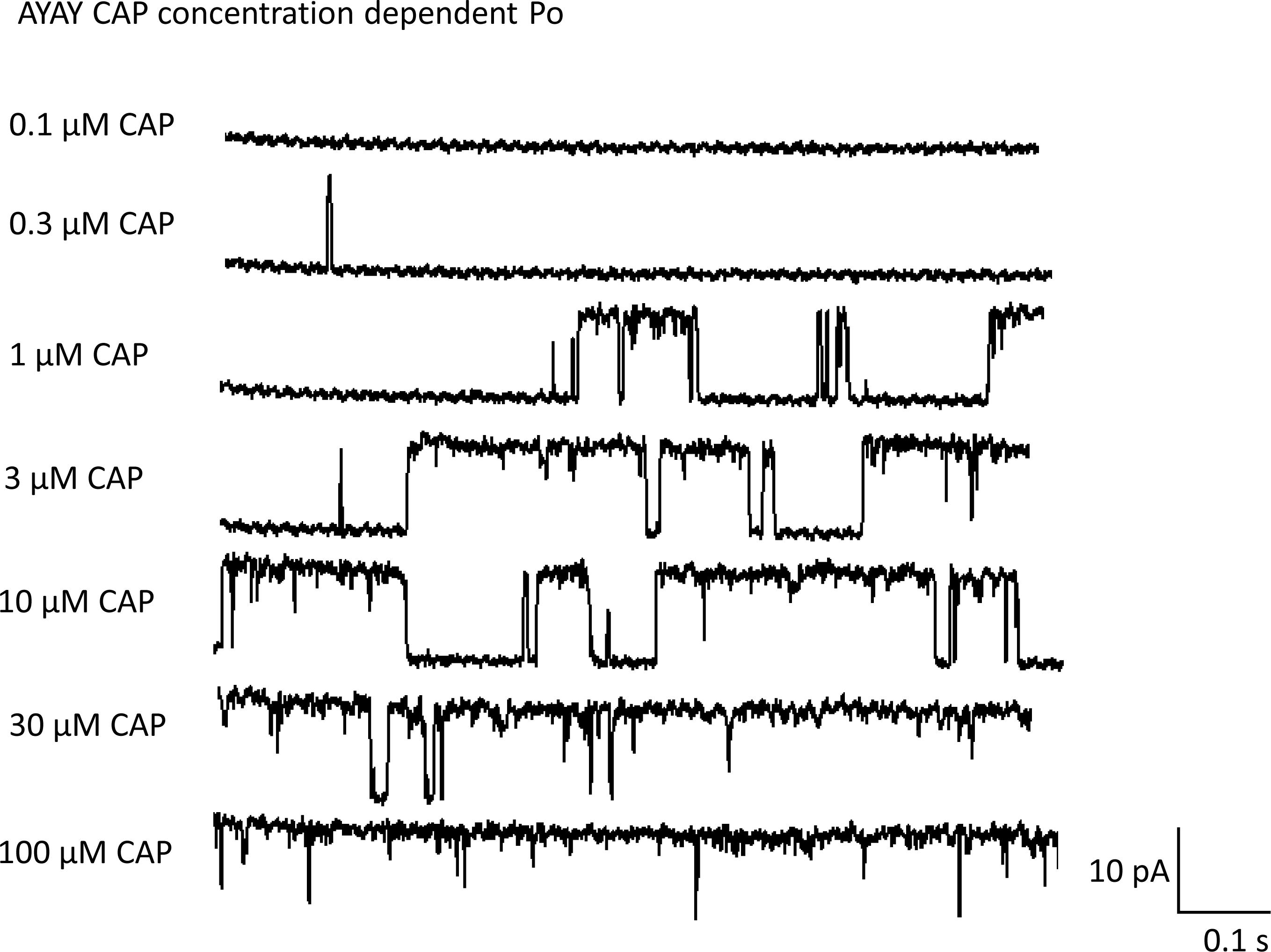
Representative single channel current traces of YAYA and AYAY concatemers activated by different concentration of capsaicin from the same inside out patch.

**Supplementary Figure 2a-b.**
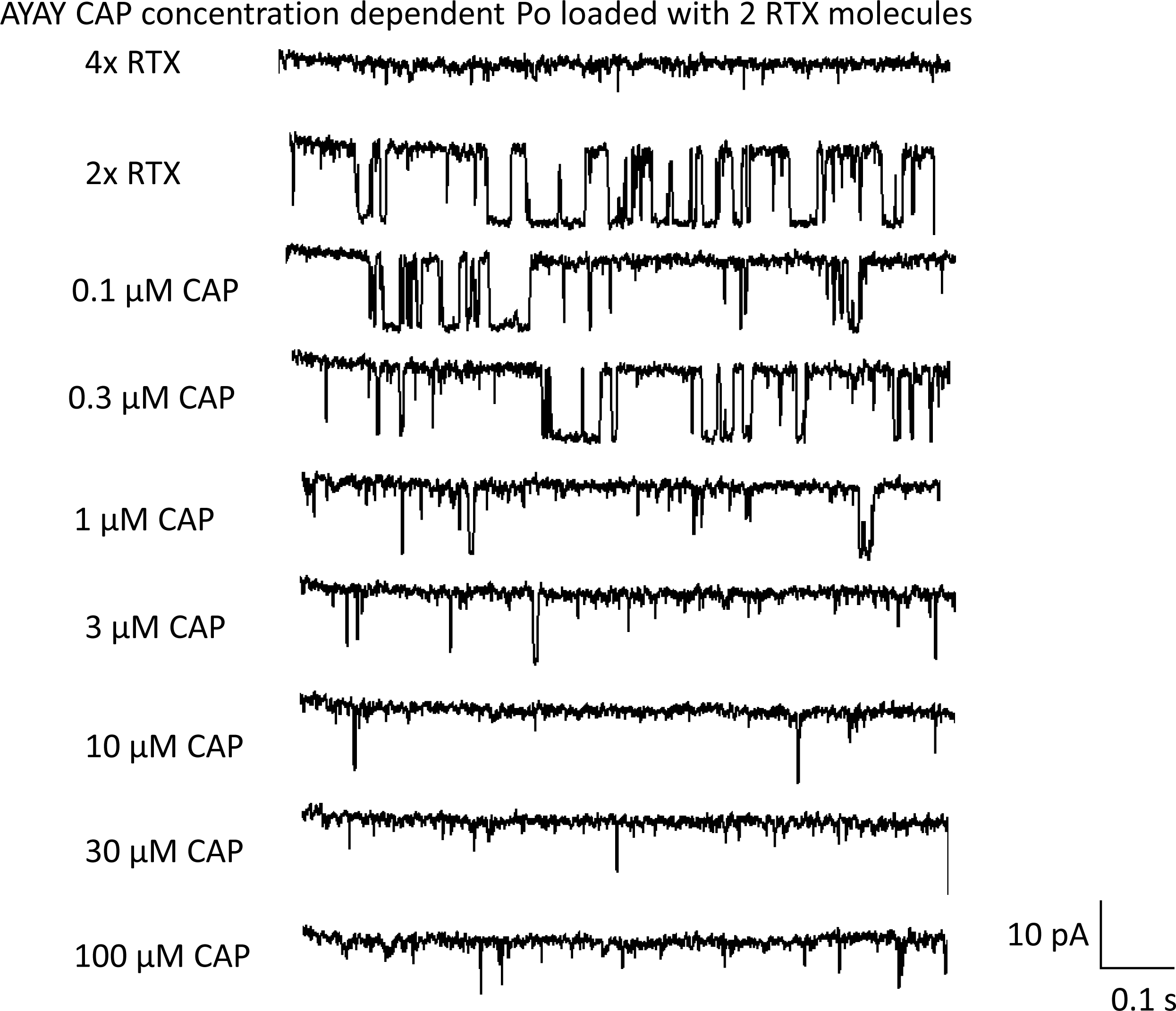
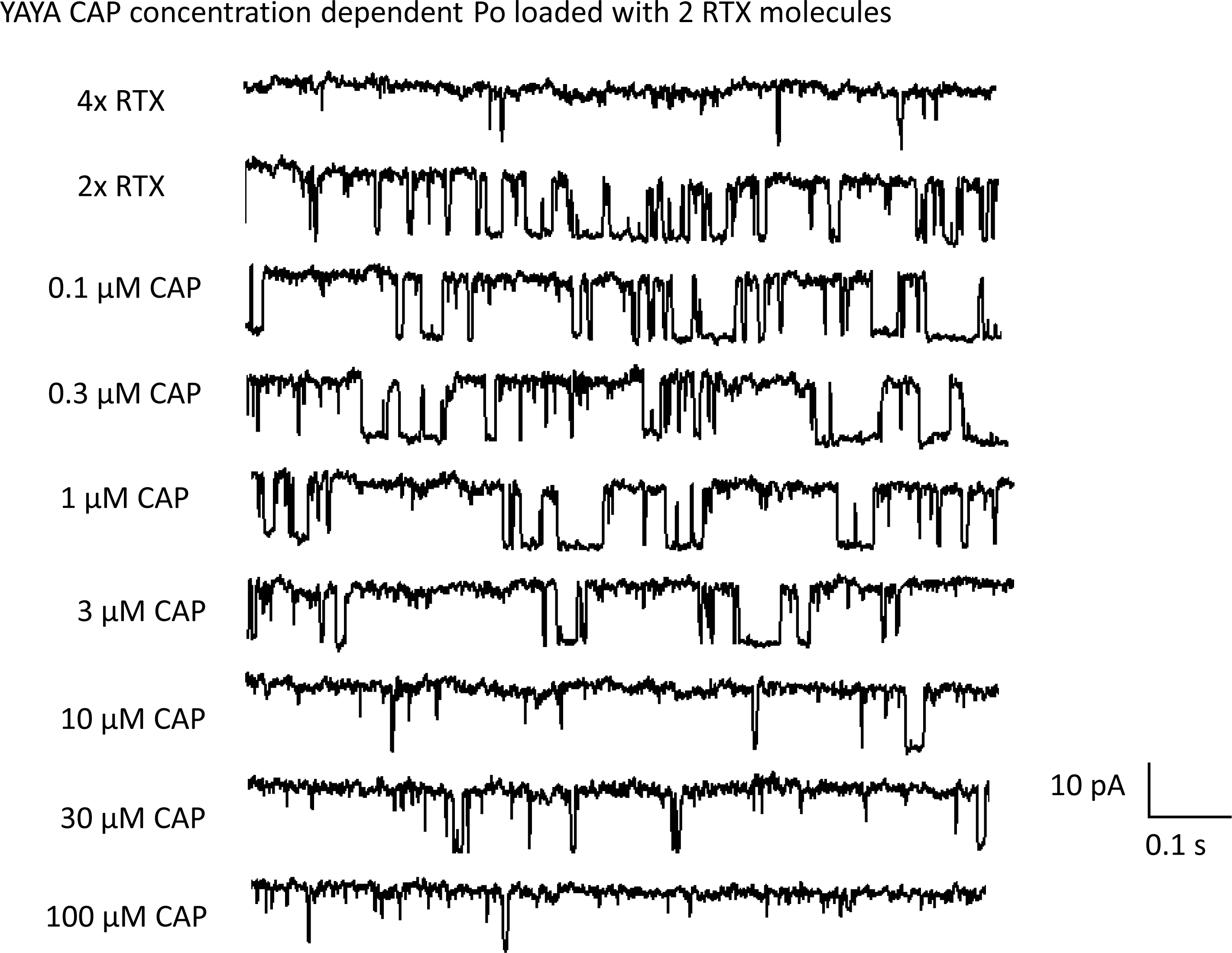
Representative single channel current traces of YAYA and AYAY concatemers first loaded with two RTX molecules in the wild type binding site then activated by different concentration of capsaicin from the same inside out patch.

**Supplementary Figure 3.**
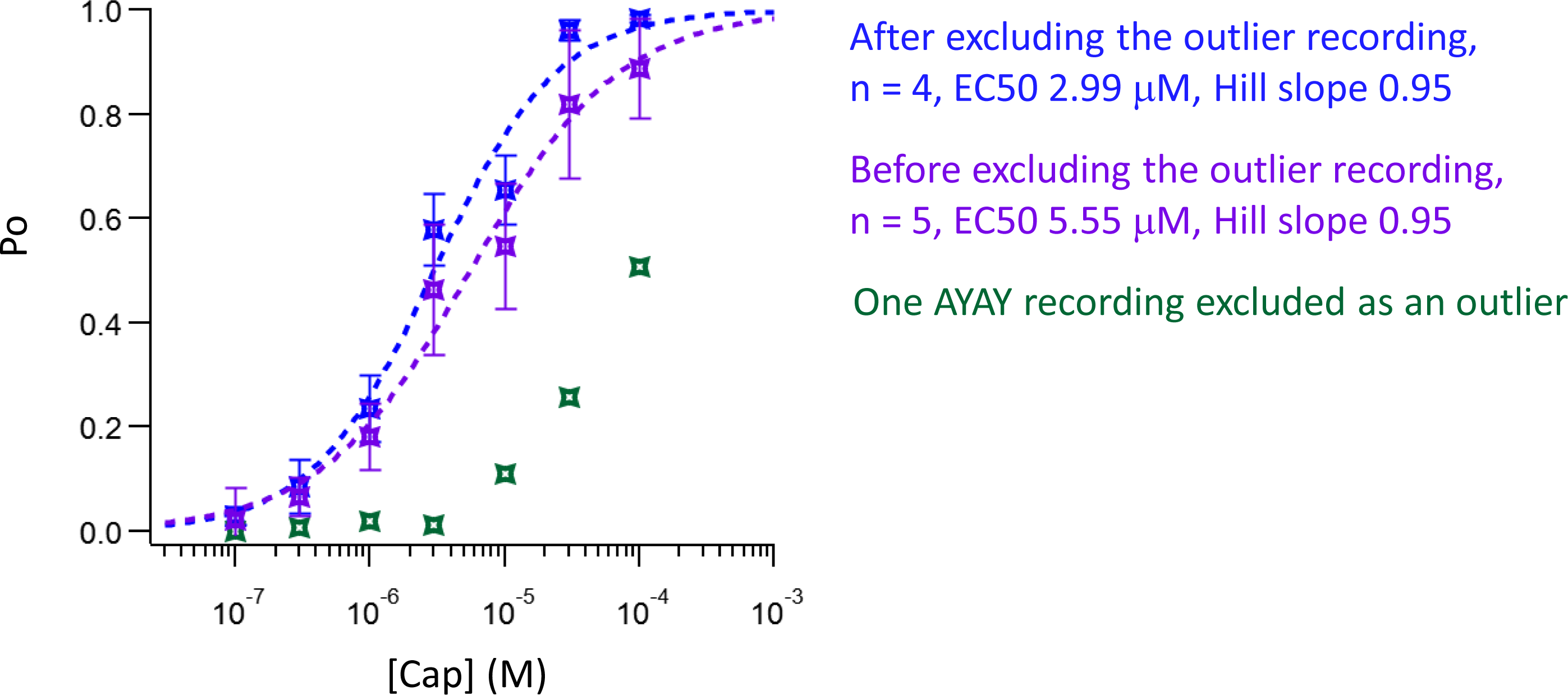
Replot of Figure 1B by including an outlier recording from AYAY channels.

## Acknowledgements

We are grateful to Avi Priel for sharing the concatemer constructs in this study. We thank current and former members of the Zheng lab for assistance and discussion. This study is supported by National Institutes of Health grant R01NS103954 (to JZ).

## Footnotes

* One of the 5 recordings from AYAY yielded an abnormal right-shifted capsaicin dependence curve, which was excluded from Figure 1B. Including this outlier recording would make the difference from YYAA even smaller (see Supplementary Figure 3).

## References

Bolton, W., and M. F. Perutz. 1970. “Three Dimensional Fourier Synthesis of Horse Deoxyhaemoglobin at 2.8 Å Resolution.” Nature 228 (5271):551–552. doi: 10.1038/228551a0.

Dong, Yawen, Yue Yin, Simon Vu, Fan Yang, Vladimir Yarov-Yarovoy, Yuhua Tian, and Jie Zheng. 2019. “A distinct structural mechanism underlies TRPV1 activation by piperine.” Biochemical and Biophysical Research Communications 516 (2):365–372. doi: 10.1016/j.bbrc.2019.06.039.

Edsall, John T. 1972. “Blood and hemoglobin: The evolution of knowledge of functional adaptation in a biochemical system.” Journal of the History of Biology 5 (2):205–257. doi: 10.1007/BF00346659.

Hazan, Adina, Rakesh Kumar, Henry Matzner, and Avi Priel. 2015. “The pain receptor TRPV1 displays agonist-dependent activation stoichiometry.” Scientific reports 5 (1):1–13.

Hodgkin, A. L., and A. F. Huxley. 1952. “A quantitative description of membrane current and its application to conduction and excitation in nerve.” J Physiol 117 (4):500–44. doi: 10.1113/jphysiol.1952.sp004764.

Horrigan, F. T., and R. W. Aldrich. 1999. “Allosteric voltage gating of potassium channels II. Mslo channel gating charge movement in the absence of Ca(2+).” J Gen Physiol 114 (2):305–36.

Horrigan, F. T., and R. W. Aldrich. 2002. “Coupling between voltage sensor activation, Ca2+ binding and channel opening in large conductance (BK) potassium channels.” J Gen Physiol 120 (3):267–305.

Horrigan, F. T., J. Cui, and R. W. Aldrich. 1999. “Allosteric voltage gating of potassium channels I. Mslo ionic currents in the absence of Ca(2+).” J Gen Physiol 114 (2):277–304.

Koshland, D. E., Jr., G. Némethy, and D. Filmer. 1966. “Comparison of experimental binding data and theoretical models in proteins containing subunits.” Biochemistry 5 (1):365–85. doi: 10.1021/bi00865a047.

Li, Shisheng, Phuong Tran Nguyen, Simon Vu, Vladimir Yarov-Yarovoy, and Jie Zheng. 2023. “Opening of capsaicin receptor TRPV1 is Stabilized Equally by Its Four Subunits.” Journal of Biological Chemistry:104828.

Li, Shisheng, and Jie Zheng. 2023. “The Capsaicin Binding Affinity of Wild-Type and Mutant TRPV1 Ion Channels.” Journal of Biological Chemistry:105268. doi: 10.1016/j.jbc.2023.105268.

Liao, Maofu, Erhu Cao, David Julius, and Yifan Cheng. 2013. “Structure of the TRPV1 ion channel determined by electron cryo-microscopy.” Nature 504 (7478):107–112.

Monod, J., J. Wyman, and J. P. Changeux. 1965. “On the Nature of Allosteric Transitions: A Plausible Model.” J Mol Biol 12:88–118. doi: 10.1016/s0022-2836(65)80285-6.

Rothberg, B. S., and K. L. Magleby. 1998. “Kinetic structure of large-conductance Ca2+-activated K+ channels suggests that the gating includes transitions through intermediate or secondary states. A mechanism for flickers.” J Gen Physiol 111 (6):751–80. doi: 10.1085/jgp.111.6.751.

Schoppa, N. E., and F. J. Sigworth. 1998. “Activation of Shaker potassium channels. III. An activation gating model for wild-type and V2 mutant channels.” J Gen Physiol 111 (2):313–42. doi: 10.1085/jgp.111.2.313.

Vu, S., V. Singh, H. Wulff, V. Yarov-Yarovoy, and J. Zheng. 2020. “New capsaicin analogs as molecular rulers to define the permissive conformation of the mouse TRPV1 ligand-binding pocket.” Elife 9. doi: 10.7554/eLife.62039.

Yang, Fan, Xian Xiao, Wei Cheng, Wei Yang, Peilin Yu, Zhenzhen Song, Vladimir Yarov-Yarovoy, and Jie Zheng. 2015. “Structural mechanism underlying capsaicin binding and activation of the TRPV1 ion channel.” Nature chemical biology 11 (7):518–524.

Yang, Fan, Xian Xiao, Bo Hyun Lee, Simon Vu, Wei Yang, Vladimir Yarov-Yarovoy, and Jie Zheng. 2018. “The conformational wave in capsaicin activation of transient receptor potential vanilloid 1 ion channel.” Nature communications 9 (1):1–9.

Yin, Yue, Yawen Dong, Simon Vu, Fan Yang, Vladimir Yarov-Yarovoy, Yuhua Tian, and Jie Zheng. 2019. “Structural mechanisms underlying activation of TRPV1 channels by pungent compounds in gingers.” British Journal of Pharmacology 176 (17):3364-3377.

Zagotta, W. N., T. Hoshi, and R. W. Aldrich. 1994. “Shaker potassium channel gating. III: Evaluation of kinetic models for activation.” J Gen Physiol 103 (2):321–62. doi: 10.1085/jgp.103.2.321.

Zhang, Kaihua, David Julius, and Yifan Cheng. 2021. “Structural snapshots of TRPV1 reveal mechanism of polymodal functionality.” Cell 184 (20):5138–5150. e12.

Zheng, Jie, and Matthew C Trudeau. 2023. Textbook of Ion Channels, CRC Press.

